# Asymmetric binding of coactivator SRC1 to FXR-RXR and allosteric communication within the complex

**DOI:** 10.1101/2024.05.13.593894

**Authors:** Yanan Sheng, Yaoting Guo, Mingze Sun, Yan Dong, Yue Yin, Yanwu Wang, Chao Peng, Yong Xu, Na Wang, Jinsong Liu

**Author notes:** These authors contributed equally (Yanan Sheng, Yaoting Guo). Correspondence (N. W.), (J. L.).

## Abstract

Farnesoid X receptor (FXR) is a promising target for treatment of metabolic associated fatty liver disease (MAFLD). In this study, we employed an integrative approach to investigate the interaction between FXR-RXRα-hSHP-1 complex and the entire coactivator SRC1-NRID (nuclear receptor interaction domain). We constructed a multi-domain model of FXR_120_-RXRα_98_-hSHP-1, highlighting the interface between FXR-DBD and LBD. Using HDX-MS, XL-MS, and biochemical assays, we revealed the allosteric communications in FXR-RXRα-hSHP-1 upon agonist and DNA binding. We then demonstrated that SRC1 binds only to the coactivator binding surface of FXR within the FXR-RXRα heterodimer, with the NR-box2 and NR-box3 of SRC1 as the key binding motifs. Our findings, which provide the first model of SRC1-NRID in complex with FXR-RXRα-hSHP-1, shed light on the molecular mechanism through which the coactivator asymmetrically interacts with nuclear receptors and provide structural basis for further understanding the function of FXR and its implications in diseases.

## Introduction

Metabolic dysfunction-associated steatohepatitis (MASH) is a chronic liver disease that progresses from metabolic associated fatty liver disease (MAFLD), becoming the main cause of liver cirrhosis and cancer(1, 2). Due to the intricate pathogenesis of MASH, only one drug has been officially approved for its treatment very recently(3). The agonist of farnesoid X receptor (FXR, NR1H4), obeticholic acid (OCA), was the first MASH drug to enter phase III clinical trials worldwide(4), as FXR being one of the most promising targets for the treatment of MASH. However, it was found that OCA treatment had serious side effects(5). Therefore, studies on the molecular mechanisms of the regulation of FXR transcription is urgently needed for designing more effective drug with less side effects.

FXR is a ligand-dependent activated type II nuclear receptor (NR) that shares a common structure with classical NRs, consisting of an N-terminal domain (NTD), a DNA-binding domain (DBD), a hinge region, and a ligand-binding domain (LBD)(6–8). Typically, FXR functions as a heterodimer alongside the retinoid X receptor (RXR)(6, 9). The FXR-DBD typically regulates gene transcription by binding to specific DNA sequences called farnesoid X receptor response elements (FXREs), which usually consist of a repeated pattern of G/AGGTCA with a one base pair in-between (IR1) (6, 10). The LBD of FXR consists of 12 α-helices, in which houses not only an LBP (ligand-binding-pocket) but also a dimerization surface and a binding site for cofactor, serving multiple functions(11–13). Various crystal structures of FXR-LBD with ligands have been resolved(14–16). We previously presented the structure of FXR-RXRα-LBD heterodimer in the presence of coactivator peptides and FXR/RXRα agonists(12). Of late, the crystal structure of FXR-RXRα-DBD and DNA has also been reported(17). However, how the individual domains work together in a multi-domain NR is still poorly understood. Studies have shown that there is extensive allosteric communication in NRs, i.e. the structural and dynamic changes between different domains affect each other to achieve the functional regulation of NRs(18–22). Therefore, structural study of the full-length FXR-RXRα protein is crucial for in-depth understanding of the interaction, allosteric communication, and signal transduction mechanisms of the multi-domain NRs.

To achieve full activity, FXR-RXRα must associate with coactivators that connect FXR-RXRα to the basal transcriptional machinery to remodel the chromatin and alter gene transcription(23–25). SRC1, a primary coactivator directly binding to FXR-RXRα and further mediating the recruitment of secondary coactivators such as p300 and CARM1, is essential for the formation of the FXR transcriptional activation complex(23, 25, 26). SRC1 binds to the LBD of FXR-RXRα via a nuclear receptor interaction domain (NRID) containing three highly conserved α-helical LXXLL motifs (NR-box) (27–31). The crystal structure from our previous study reveals that each monomer in FXR-RXRα-LBD heterodimer recruits one SRC1 peptide, containing one LXXLL motif, to the corresponding coactivator binding surface^12^. The leucine residues from the LXXLL motif are embedded in a hydrophobic groove in LBD and locked by a charge clamp composed of lysine (K) from H3 and glutamate (E) from H12 (12, 32, 33). However, all crystal structures reported up to date only contain a single LXXLL motif peptide due to the disordered feature in SRC1-NRID(34). Recently, small molecule inhibitors have been developed targeting SRC3 and SRC1 to disrupt their interactions with NRs, resulting in inhibition of their transcriptional activation functions and further anti-tumor activity(35–37).

In this study, we used integrative approaches to investigate the structure and function of the complex between FXR-RXRα and SRC1. We constructed the first multi-domain model of the FXR-RXRα with an IR1 sequence and revealed the interaction between the DBD and LBD of FXR. Furthermore, we revealed an asymmetric recruitment of SRC1 by FXR-RXRα for the first time. Through identifying NR-box2 and NR-box3 of the SRC1-NRID as key players in binding to FXR-RXRα, we put forth a structure model for the complex of SRC1_630-987_ with FXR_120_-RXRα_98_-hSHP-1. These results will greatly advance our understanding on the molecular organization and allosteric communication within the FXR transcriptional activation complex and offer new insights for the development of FXR-related drugs.

## Results

### FXRE and ligands co-regulate SRC1 binding to FXR-RXRα

To investigate the effects of FXRE and agonists on SRC1 binding to FXR-RXRα, we expressed and purified SRC1_630-987_ and FXR_120_-RXRα_98_. SRC1_630-987_ includes an NRID containing three NR-boxes and a p300-interacting domain (Fig. 1A). The DNA used in this study was the FXRE (IR1) locating in the promoter region of the human small heterodimer chaperone 1 (hSHP-1) gene(38) (Fig. 1B). Biofilm interferometry (BLI) was employed to assess the binding affinity of SRC1_630-987_ for various states of FXR_120_-RXRα_98_.

**Figure 1.**
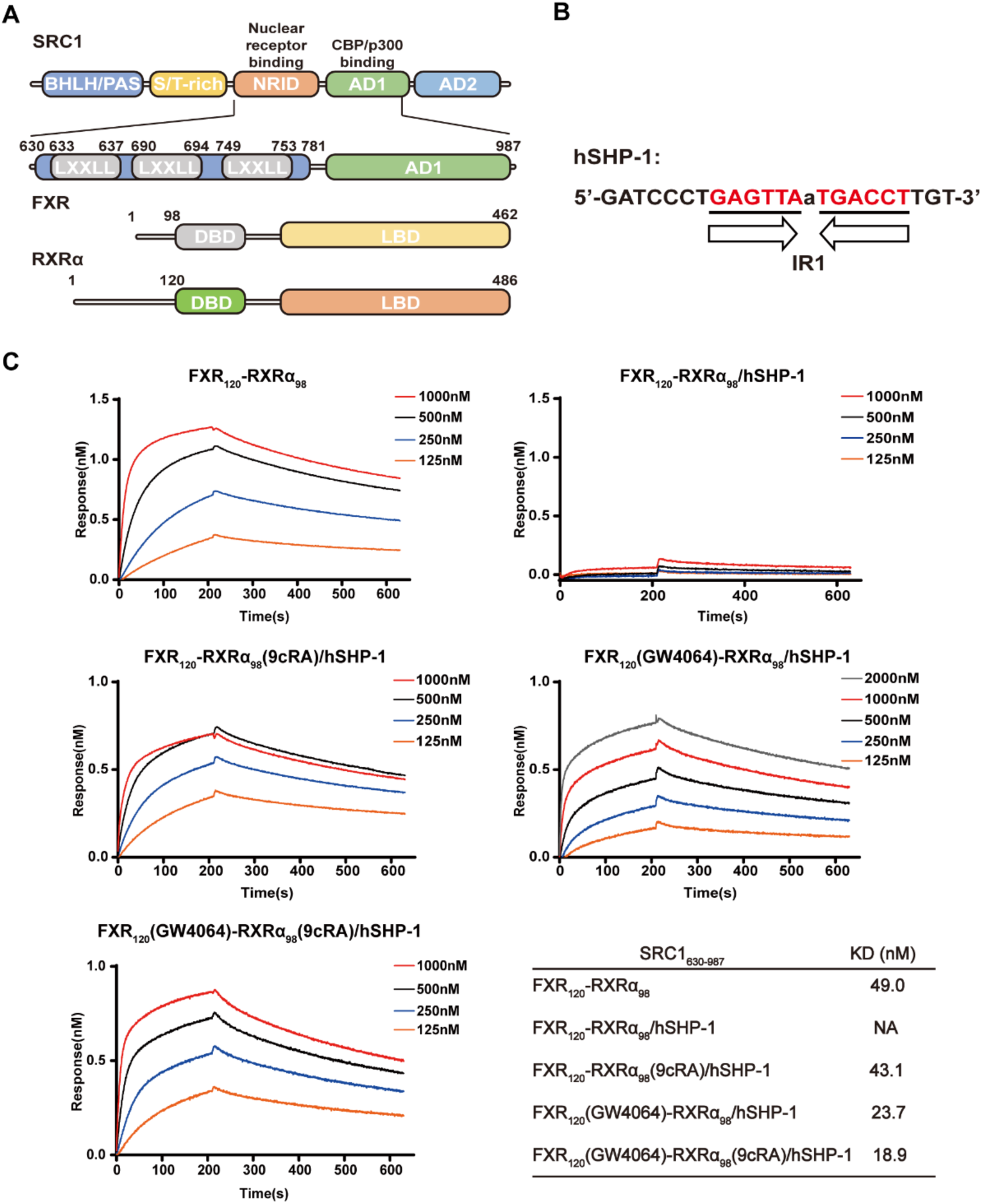
FXRE and ligand co-regulate SRC1 binding to FXR-RXRα. (A) Schematic diagram of the domain architecture of SRC1, FXR and RXRα. Truncations utilized in the experiments are indicated. (B) The FXR response element in the promoter of the FXR target gene hSHP-1 used in the experiment, comprising AGGTCA inverted repeats separated by a nucleotide. (C) Binding profiles of different states of FXR120-RXRα98 to SRC1630-987 in a concentration gradient. Biotin-labeled SRC1630-987 was immobilized on the sensor, and the effects of FXR and RXRα agonists, as well as FXRE (hSHP-1), on its binding affinity to FXR120-RXRα98, were examined. The KD values are also presented in the table.

The binding experiments showed that FXR_120_-RXRα_98_ binds to SRC1_630-987_ with a KD value of 49.0 nM. However, when FXR_120_-RXRα_98_ binds to hSHP-1, it fails to bind with SRC1_630-987_ without an agonist (Fig. 1C). This suggests that the presence of hSHP-1 alters the conformation of the LBD of FXR_120_-RXRα_98_, preventing its binding to SRC1_630-987_. Furthermore, when the RXRα agonist 9cRA (9-cis-Retinoic Acid) is present, FXR_120_-RXRα_98_-hSHP-1 can bind to SRC1_630-987_ with a KD value of 43.1 nM (Fig. 1C). This indicates that agonist binding changes the conformation of FXR_120_-RXRα_98_, enabling it to recruit SRC1_630-987_. Remarkably, in the presence of the FXR agonist GW4064, FXR_120_-RXRα_98_-hSHP-1 binds to SRC1_630-987_ at a KD value of 23.7 nM, demonstrating a stronger effect for FXR agonist binding (Fig. 1C). These findings suggest that both hSHP-1 and agonists influence the binding of SRC1_630-987_ to FXR_120_-RXRα_98_. Additionally, we also found that further addition of 9cRA in the presence of GW4064 only slightly increases the affinity of SRC1_630-987_ for FXR_120_-RXRα_98_, indicating that FXR may play a dominant role in recruiting SRC1 (Fig. 1C). These results demonstrate that the binding of SRC1_630-987_ to FXR_120_-RXRα_98_ is co-regulated by FXRE and ligands.

### Integrative modelling of the FXR-RXRα heterodimer binding to hSHP-1

To gain insight into the molecular mechanisms by which FXRE and agonists co-regulate the binding of SRC1 to FXR-RXRα, we constructed an FXR-RXRα-hSHP-1 model by integrating various biophysical approaches. Bis (sulfosuccinimidyl) suberate (BS3) and 1-ethyl-3-[3-dimethylaminopropyl] carbodiimide hydrochloride (EDC) were used to cross-link FXR_120_-RXRα_98_-hSHP-1, respectively. The complex band at 80 kDa was excised for mass spectrometry analysis (Fig. S1A). Both BS3 and EDC produced similar cross-linking results (Fig. S1B and C). The multi-domain model of FXR_120_-RXRα_98_-hSHP-1 was constructed by docking the crystal structures of FXR-RXRα-LBD (PDB ID: 5Z12) and FXR-RXRα-DBD-IR1 (PDB ID:8HDM) using the LZerD web server(39, 40), together with distance constraints obtained from XL-MS (Fig. 2A). We also collected SAXS data of FXR_120_-RXRα_98_-hSHP-1. The theoretical profile calculated based on the model of FXR_120_-RXRα_98_-hSHP-1 is consistent with the experimental SAXS profile, with a χ^2^ value of 2.76 (Fig. S1F). The FXR-RXRα-hSHP-1 model also fits well with the ab initio envelope (Fig. S1G).

**Figure 2.**
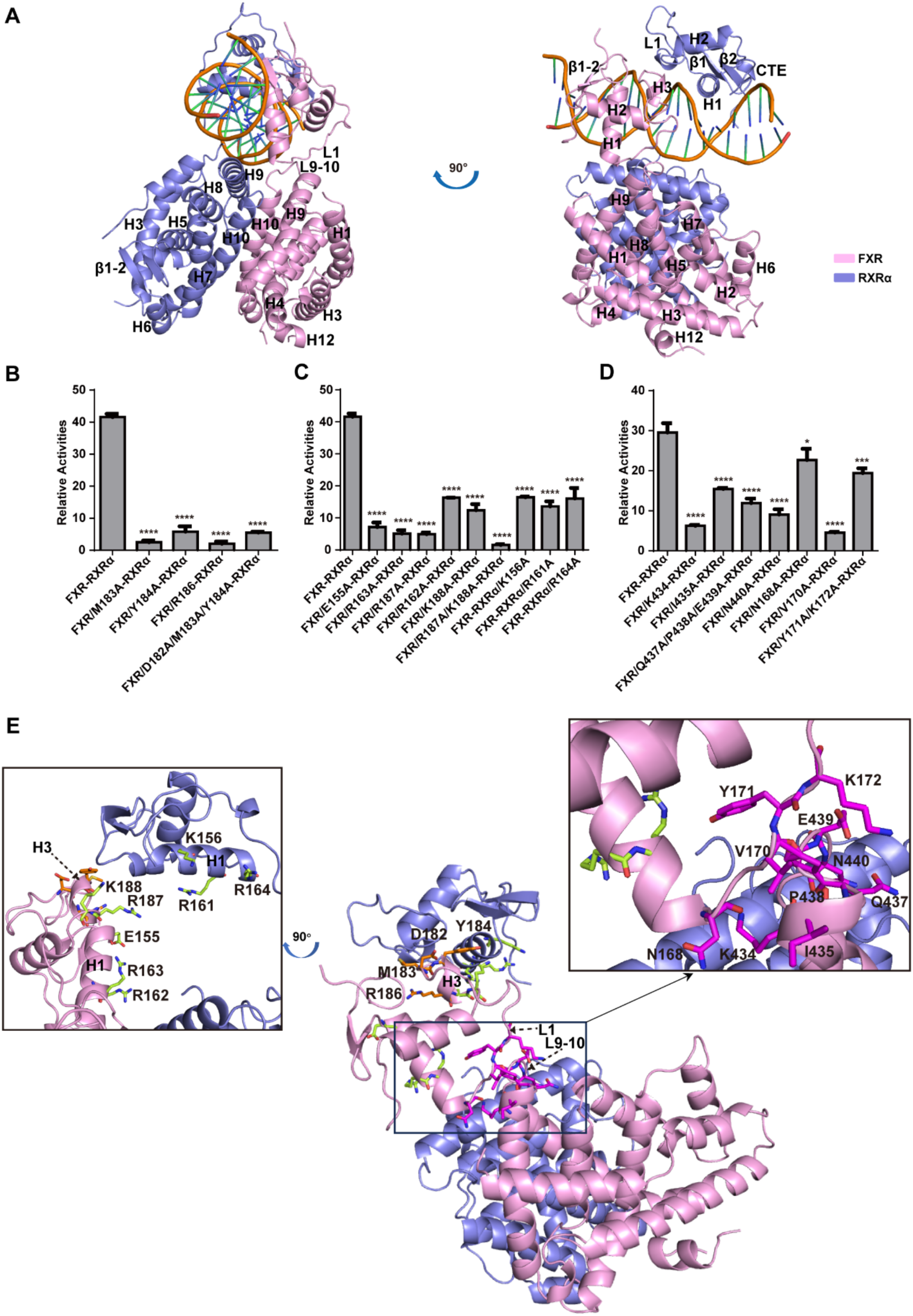
Integrative modelling of the FXR-RXRα heterodimer binding to hSHP-1. (A) Multi-domain model of FXR_120_-RXRα_98_-hSHP-1 (FXR, pink; RXRα, slate). (B) The impact of FXR mutations at the FXR-DBD and RXRα-DBD interface. (C) Transcriptional activity resulting from mutations in FXR and RXRα at the putative DNA binding interface of FXR-RXRα. (D) Transcriptional activity resulting from mutations in FXR at the FXR-LBD and FXR-DBD interface. (E) The mutated residues mapped to the FXR-RXRα-hSHP-1 model. The amino acids mutated at the interface between FXR-DBD and RXRα-DBD are highlighted in orange. The lime color represents the mutant residues at the DNA-binding interface of FXR-RXRα-DBD. The magenta color indicates the mutated residues located at the interface between FXR-LBD and FXR-DBD.

To assess the impact of domain interfaces in the FXR_120_-RXRα_98_-hSHP-1 model on the transcriptional activity, cellular functional experiments were conducted. The cross-linking data of FXR_120_-RXRα_98_-hSHP-1 were consistent with the crystal structure of FXR-RXRα-LBD (Fig. S1D). Therefore, no mutation experiments were performed on the dimerization interface between LBDs. Whereas the cross-link data did not agree completely with the crystal structure of FXR-RXRα-DBD-IR1 (Fig. S1E). As a result, specific amino acids on FXR (M183A, Y184A, D182A/M183A/Y184A) located at the interface between FXR-DBD and RXRα-DBD were mutated. (Fig. 2E). Mutations at these sites resulted in a significant decrease in the transcriptional activity of FXR-RXRα on hSHP-1 (Fig. 2B). We also generated the mutations on the residues in H1, which is inserted into the major groove of DNA, on the DBD of FXR (E155A, R162A, R163A) and RXRα (K156A, R161A, R164A) (Fig. 2E). Results showed that these residues are critical for the binding of FXR-RXRα-DBD to DNA (Fig. 2C). Moreover, we mutated the residues in H3 of the FXR-DBD in the model (R187A, R188A, R187A/R188A) that is shown to have contact with the DNA minor groove (Fig. 2E). These mutations also significantly reduced the transcriptional activity of FXR-RXRα on hSHP-1 (Fig. 2C). Collectively, the results demonstrated that the binding of FXR-RXRα-DBD to hSHP-1 in the FXR_120_-RXRα_98_-hSHP-1 model agrees with those in the FXR-RXRα-DBD-IR1 crystal structure.

More important, there is also an interface between the FXR-LBD and DBD in our model. Specifically, the interface is between L9-10 (H9-H10 loop) of FXR-LBD and L1 (the loop following H1) of FXR-DBD (Fig. 2E). We designed mutations on the critical residues (K434A, I435A, Q437A/P438A/E439A, F440A of FXR-LBD; N168A, V170A, Y171A/K172A of FXR-DBD) (Fig. 2F). As shown in Figure 2D, mutating these amino acids significantly reduced the transcriptional activity of FXR-RXRα on hSHP-1.

Altogether, these results demonstrated that these three interfaces presented in the model are important for the transcriptional activity of the FXR-RXRα.

### Allosteric communication between the LBD and DBD of FXR-RXRα

To investigate the molecular mechanism by which FXRE and agonists influence the recruitment of SRC1 by FXR-RXRα, we utilized HDX-MS to examine the allosteric communication between LBD and DBD of FXR-RXRα induced by agonists and DNA binding (Fig. 3A). By integrating the HDX data with the FXR_120_-RXRα_98_-hSHP-1 model, we were able to gain a more comprehensive understanding of the HDX results (Fig. 3F and G).

**Figure 3.**
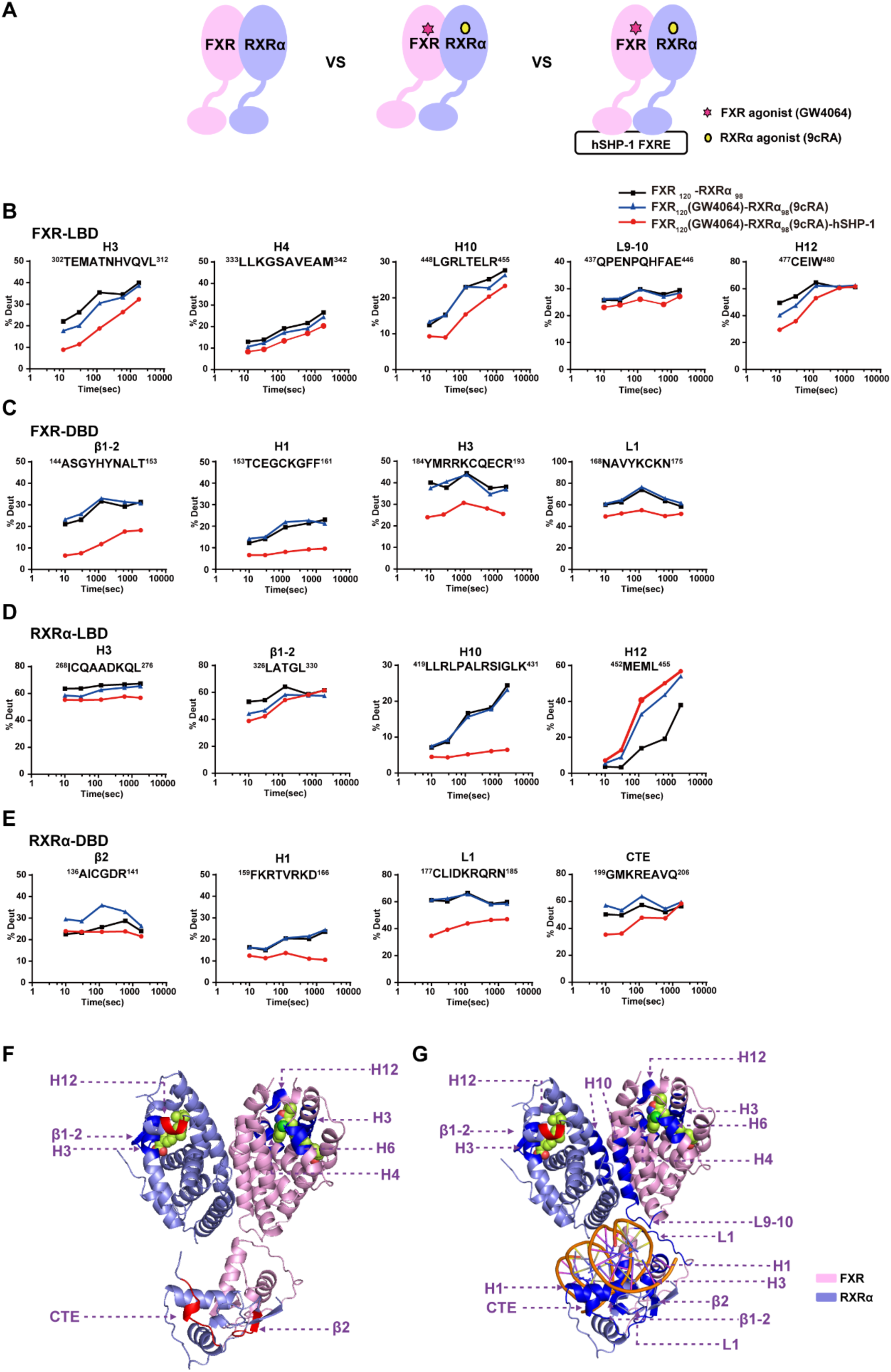
Allosteric communication between the LBD and DBD of FXR-RXRα. (A) Schematic diagram of the HDX-MS experiment. Plots of deuterium uptake by FXR (B and C) and RXRα (D and E) as a function of deuteration time after binding to agonists and DNA. (F) Comparison of FXR-RXRα and FXR120(GW4064)-RXRα98(9cRA) as well as (G) FXR120(GW4064)-RXRα98(9cRA) and FXR120(GW4064)-RXRα98(9cRA)-DNA peptides with reduced deuterium uptake (dark blue) and peptides with elevated deuterium (red) in HDX-MS were mapped to the FXR120-RXRα98 model (FXR, pink; RXRα, slate; GW4064 in FXR and 9cRA in RXRα are shown in space-filling representation colored by atom type: oxygen as red, nitrogen as blue, sulfur as yellow, chlorine as green, and carbon as pink.).

First, we studied the conformational changes of FXR_120_-RXRα_98_ upon binding with their agonists. Comparing the HDX results of FXR_120_-RXRα_98_ and FXR_120_(GW4064)-RXRα_98_(9cRA) (Fig. 3B, C, D and E) demonstrated that the regions in FXR-RXRα-LBD with reduced solvent exposure upon ligand binding are comparable to those directly interact with the ligand in the crystal structure. Specifically, regions such as H3, H4, H12 (Fig. 3B and F), and H6 (Fig. S2C) in FXR-LBD, as well as H3 and β1-2 in RXRα-LBD, exhibited significant reductions in HDX (Fig. 3B, D and F). Agonists binding stabilizes the LBP of FXR and subsequently creates a surface that facilitates the recruitment of coactivators, as shown in Figure 3F. Surprisingly, upon agonist binding, H12 within the RXRα-LBD exhibits an accelerated rate of HDX (Fig. 3D and F), suggesting that agonist binding increases the dynamics and flexibility of H12 within the RXRα-LBD. The DBD region of FXR_120_-RXRα_98_ also exhibits obvious HDX changes. As shown in Figure 3E and F, the HDX rates of the N-terminal β2 and CTE of RXRα-DBD are significantly higher, suggestive of increased flexibility in these regions. Apparently, ligand binding, even in the absence of direct contact with the DBD, can induce an allosteric effect that is propagated to the distal end of FXR_120_-RXRα_98_.

Then, we investigated the impact of FXR_120_-RXRα_98_ conformational dynamics upon its interaction with hSHP-1, in the presence of agonists such as GW4064 and 9cRA. Comparison of the HDX between FXR_120_(GW4064)-RXRα_98_(9cRA) and FXR_120_(GW4064)-RXRα_98_(9cRA)-hSHP-1 revealed that the DBD, LBD, and hinge region (Fig. S2C) of both FXR and RXRα underwent significant HDX changes. Upon DNA binding, there is a reduction in deuterium uptake in H3, H4, H6 (Fig. S2C), and H12 of the FXR-LBD, as well as in H3 and the β1-2 of the RXRα-LBD (Fig. 3B, D, and G). These findings suggest that DNA binding enhances the stability of the coactivator binding surface in FXR, which can facilitate the coactivator recruitment. Furthermore, H10 of both the FXR-LBD and RXRα-LBD, which are located at the FXR-RXRα-LBD dimerization interface, exhibit a significant decrease in HDX upon DNA binding (Fig. 3B, D, andG). Additionally, our XL-MS results (Fig. S2A and B) show that DNA binding significantly increases the number of cross-links between the LBDs of the FXR and RXRα. These findings suggest that DNA binding stabilize the dimerization between the FXR-LBD and RXRα-LBD. And it has been reported that the dimerization of FXR-RXRα-LBD increases its affinity for SRC1(12). Our data indicate that the binding of DNA may promote the recruitment of SRC1 to FXR-RXRα-LBD via the dimerization of FXR-RXRα-LBD.

In the DBDs, the HDX reduction observed in H1 of both FXR-DBD and RXRα-DBD (Fig. 3C, E, and G) is consistent with their insertion into the major groove of DNA. When compared to RXRα-DBD, H1 in FXR-DBD exhibits stronger solvent exchange protection, suggesting a stronger interaction with DNA compared to RXRα (Fig. 3C and E). The H3 in FXR-DBD and L1 (the loop following H1) of RXRα-DBD peptides also exhibited significant solution protection (Fig. 3C and E), further confirming the interface between FXR-DBD and RXRα-DBD as presented in the model (Fig. 3G). Furthermore, the deuterium uptake of the N-terminal β1-2 of FXR-DBD was significantly reduced, indicating its involvement in DNA binding (Fig. 3C and G). Additionally, mutating key amino acids in this region led to a significant reduction in FXR-RXRα transcriptional activity (Fig. S2D). Overall, besides the recognition helix of DBD, the NTDs of both FXR and RXRα participate in DNA binding and play a crucial role in transcriptional activation of target genes by FXR-RXRα. Interestingly, the L9-10 of FXR-LBD and L1 of FXR-DBD, locating at the interface between FXR-LBD and DBD, showed reduced deuterium uptake (Fig. 3B, C, and G). This strongly supports the interaction between FXR-DBD and FXR-LBD (Fig. 2B). Moreover, significant solution protection was observed in the hinge region of FXR (Fig. S2C). And our XL-MS results indicated a significant increase in the number of cross-linking pairs between the DBD and LBD of FXR upon binding of hSHP-1 to FXR_120_(GW4064)-RXRα_98_(9cRA) (Fig. S2A and B). These results suggest that the distance between FXR-DBD and LBD may become closer. Based on these findings, we speculate that binding of hSHP-1 induces conformational changes in FXR, leading to signal transmission between the DBD and LBD of FXR-RXRα through the dynamic allostery of L9-10 of FXR-LBD and FXR-DBD. In summary, these results suggest that DNA can facilitate allosteric communication and stabilize the coactivator binding surface of FXR with the help of agonists, leading to enhanced recruitment of coactivators such as SRC.

### Binding of SRC1 to FXR asymmetrically in the transcriptional activation of FXR-RXRα

We performed a transient transfection assay using a reporter gene under the control of three copies of FXRE (3×hSHP-1-luc) (Fig. 1A) and demonstrated that SRC1 significantly increased the activity of the reporter gene with agonists (Fig. S3A).

Then we purified the proteins SRC1_630-987_ comprising the entire NRID of SRC1 (Fig. 1A) and FXR_120_-RXRα_98_. In vitro, we assembled the SRC1_630-987_-FXR_120_-RXRα_98_-hSHP-1 complex. The formation of FXR_120_-RXRα_98_ and SRC1_630-987_ complex was confirmed by both size exclusion chromatographic analysis (SEC) (Fig. S3B) and SDS-PAGE analysis (Fig. S3C). To accurately determine the stoichiometric ratio within the complex, we employed SEC coupled with Multi-Angle Light Scattering (SEC-MALS) to measure the molecular weight of SRC1_630-987_-FXR_120_-RXRα_98_-hSHP-1. The SEC-MALS analysis demonstrated that the measured molecular weight of the SRC1_630-987_-FXR_120_-RXRα_98_-hSHP-1 complex closely matched the theoretical molecular weight of the complex formed by SRC1_630-987_ bound to FXR_120_-RXRα_98_ at a 1:1 stoichiometric ratio (Fig. 4A). These findings strongly lead to the conclusion that the ligand activated FXR_120_-RXRα_98_ heterodimer recruits only one molecule of SRC1.

**Figure 4.**
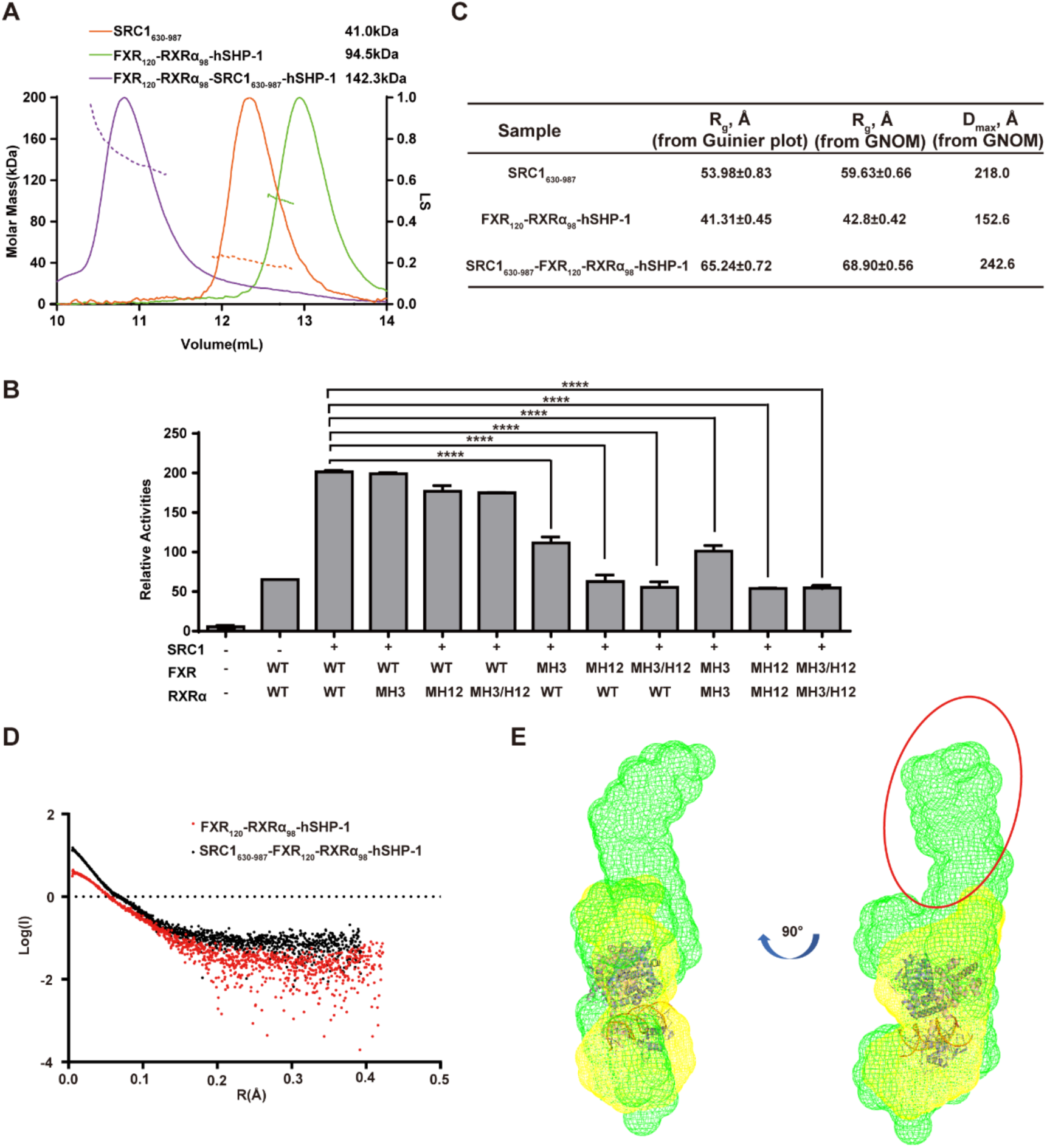
Binding of SRC1 to FXR asymmetrically in the transcriptional activation of FXR-RXRα. (A) SEC-MALS of SRC1_630-987_, FXR120-RXRα98-hSHP-1, and SRC1630-987-FXR120-RXRα98-hSHP-1 complexes. The figure shows their elution profiles on Superdex 200 10/300 GL (GE Healthcare) and the directly measured molar mass of each elution peak. (B) Luciferase reporter gene assay tests the transcriptional ability of different charge-clamp mutants for FXR and RXRα in the presence of GW4064 and 9cRA for hSHP-1. (C) SAXS parameters. R_g_ and D_max_ as determined from Guinier plot or p(r) distribution. (D) The Guinier plot for SAXS analysis of FXR120-RXRα98-hSHP-1 and SRC1630-987-FXR120-RXRα98-hSHP-1. (E) Overlay of the ab initio envelope of FXR120-RXRα98-hSHP-1 (yellow mesh) with the SRC1630-987-FXR120-RXRα98-hSHP-1 complex (green mesh) after fitting with cartoon model of FXR120-RXRα98-hSHP-1 (FXR, pink; RXRα, slate). The part circled in red is the extra electron density.

To further investigate the binding mode of SRC1 to the FXR_120_-RXRα_98_ heterodimer, mutations were introduced in the conserved coactivator binding charge clamp residues: lysine (K) residue in H3 of FXR and RXRα (mH3 mutant) and the glutamate (E) residue in H12/AF2 (mH12 mutant) (Fig. S3D and E). These mutated forms of FXR and RXRα were overexpressed in HEK293T cells. The transcriptional activity of the mutated NRs upon SRC1 stimulation was assessed by co-expression of SRC1 (Fig. 4B). Interestingly, mutations in the charge clamp on H3 of RXRα did not have any impact on the transcriptional activity of FXR-RXRα stimulated by SRC1. However, mutation of H12 in RXRα resulted in a slight reduction in transcriptional activity, possibly through weakening the agonist binding to RXRα. Whereas, mutations in the charge clamp on either H3 or H12 of FXR had a significant effect on SRC1 enhanced transcriptional activity of FXR-RXRα. Most importantly, the double mutation of the FXR charge clamp fully reduced the level of transcriptional activation of FXR-RXRα to that observed without SRC1, demonstrating the critical role of the FXR charge clamp in mediating the interaction with SRC1. Additionally, HDX-MS data showed that FXR is more protected on its coactivator binding surface when bound to DNA and agonists, which would be more favorable for its binding to SRC1 (Fig. 3B and D). Conversely, H12 of RXRα-LBD displayed increased flexibility after agonist and DNA binding (Fig. 3D), potentially suggesting a more dynamic feature for RXRα-LBD in the context of the SRC1 interaction with the heterodimer. Collectively, these results strongly suggest that SRC1 enhanced transcriptional activation of FXR-RXRα primarily results from the binding of SRC1 to FXR, not RXRα.

To delve deeper into the overall organization of the SRC1_630-987_-FXR_120_-RXRα_98_-hSHP-1 complex, SAXS data (Table S1) were collected for SRC1_630-987_, FXR_120_-RXRα_98_-hSHP-1, and SRC1_630-987_-FXR_120_-RXRα_98_-hSHP-1 complex using SEC-SAXS technique. The SAXS patterns of FXR_120_-RXRα_98_-hSHP-1 and SRC1_630-987_-FXR_120_-RXRα_98_-hSHP-1 are shown in Figure 4D, and the structural parameters are shown in Figure 4C. The ab initio structure envelope of FXR_120_-RXRα_98_-hSHP-1 after fitting with FXR_120_-RXRα_98_-hSHP-1 model was overlaid with the ab initio structure envelope of SRC1_630-987_-FXR_120_-RXRα_98_-hSHP-1 complex. There is an additional electron density at the top of FXR, which is just enough to accommodate one SRC1_630-987_ molecule, further confirming that one SRC1_630-987_ is asymmetrically bound to FXR_120_-RXRα_98_-hSHP-1 (Fig. 4E).

Collectively, these results indicate that one SRC1 molecule binds asymmetrically to the coactivator binding surface of FXR in FXR-RXRα complex via its NRID domain.

### NR-box2 and box3 of the SRC1-NRID interact with FXR-RXRα heterodimer

To identify which NR-box of SRC1 is directly involved in the interaction with FXR-RXRα, we generated various mutants in the NRID domain, including single (M1, M2, M3), double (M1/2, M2/3, M1/3), or triple (M1/2/3) NR-box mutations (Fig. 5A).

**Figure 5.**
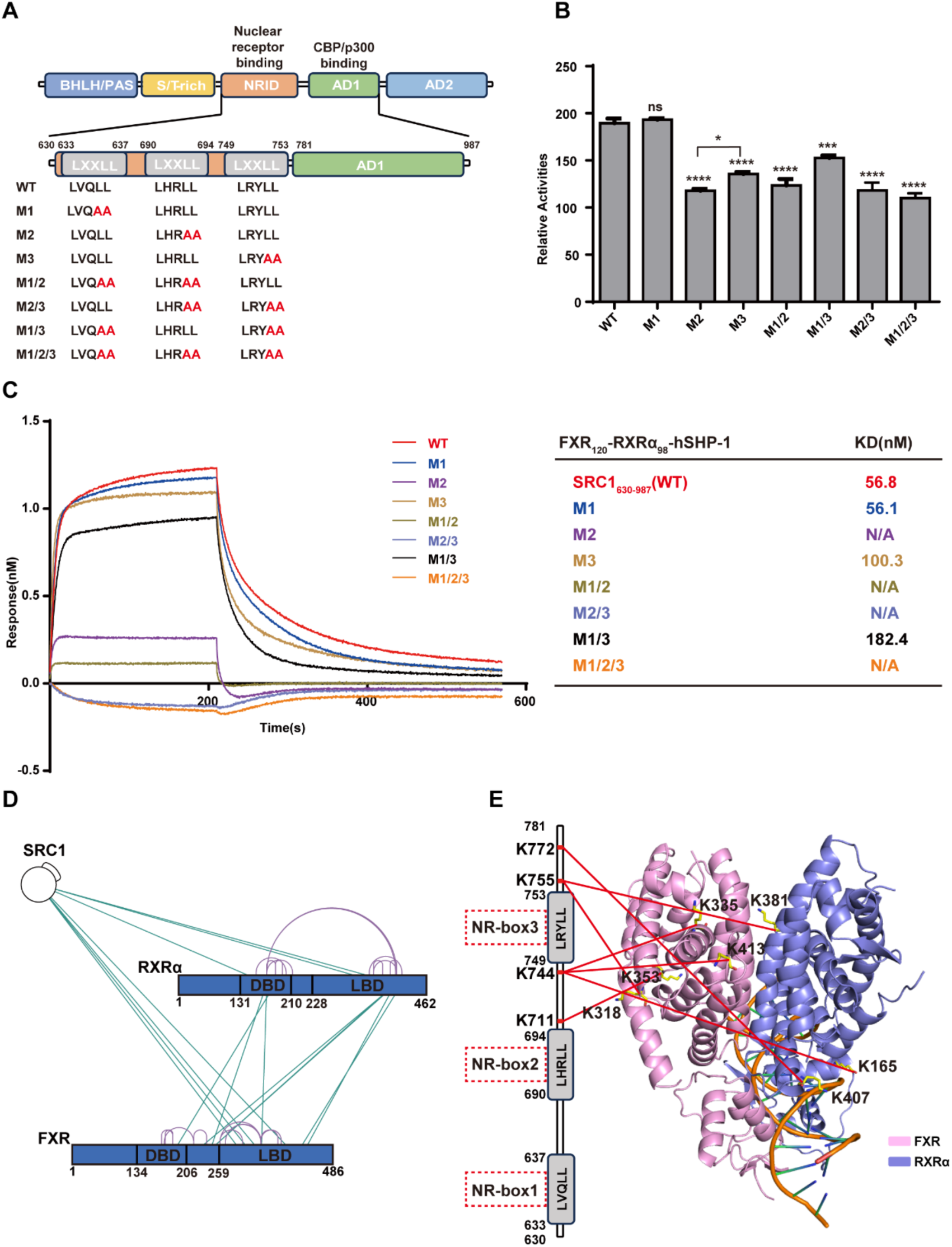
NR-box2 and box3 of the SRC1-NRID domain interact with FXR-RXRα heterodimer. (A) Schematic diagram of mutations for different NR-boxes of SRC1 used in this study. (B) Luciferase reporter assay was used to test the enhance effect of different mutants of the NR-box of SRC1 on FXR-RXRα promote hSHP-1 transcription in the presence of GW4064 and 9cRA. (C) In the presence of GW4064 and 9cRA, BLI experiments were conducted to investigate the interaction affinity of biotinylated SRC1630-987 and various NR-box mutants of SRC1630-987 with FXR120-RXRα98-hSHP-1. The calculated KD value is indicated on the right side of the graph. (D) High confidence (score ≥10) BS3 cross-linked residue maps shown on the sequence schematic of FXR and RXRα, with green lines representing cross-links and purple representing self-links. (E) Cross-linking between the NRID domain of SRC1 and FXR-RXRα is depicted in our model. The left side of the figure shows a schematic representation of the three NR-boxes in the NRID domain of SRC1, indicating the specific amino acids involved in cross-linking with FXR and RXRα. On the right-hand side, the model displays the amino acids connected to the corresponding amino acids of SRC1 that form the cross-links on the left-hand side, represented by a red line (FXR, pink; RXRα, slate).

We performed BLI assay in vitro to test the binding affinity of the SRC1_630-987_ with NR-box mutants to FXR_120_-RXRα_98_ (Fig. 5C). The results revealed that the M2 single mutant completely abolished the binding ability of SRC1_630-987_ to FXR_120_-RXRα_98_. Similarly, the M1/2, M2/3, and M1/2/3 mutants also showed a complete loss of binding ability to FXR_120_-RXRα_98_. The M3 mutant exhibited an approximately two-fold reduction in its affinity to FXR_120_-RXRα_98_. The M1/3 double mutant displayed a further decrease in affinity compared to the M3 single mutant. Interestingly, the M1 single mutant did not show significant affinity changes compared to the wild type SRC1_630-987_.These findings demonstrate that NR-box2 of the NRID domain of SRC1 plays a crucial role in binding to FXR_120_-RXRα_98_, directly interacting with the complex. Additionally, NR-box3 is also important for the interaction, potentially assisting the binding of NR-box2 to FXR_120_-RXRα_98_.

We then employed a luciferase reporter gene system to assess the effects of SRC1 mutants on FXR-RXRα transcriptional activity in cell (Fig. 5B). The results demonstrated that the M1 mutant of SRC1 did not show a significant difference in their ability to enhance FXR-RXRα transcriptional activation compared to the wild type SRC1. However, both the M2 and M3 mutants led to a notable decrease in FXR-RXRα transcriptional activation. In agreement with the binding assay, the M2 mutant exhibited a more pronounced reduction in FXR-RXRα transcriptional activity compared to the other mutants. These findings highlight the importance of NR-box2 and NR-box3 within the NRID of SRC1 for the transcriptional activation function of FXR-RXRα.

To gain further insights into the specific binding details between SRC1_630-987_ and FXR_120_-RXRα_98_, we utilized XL-MS experiments with BS3 to analyze the SRC1_630-987_-FXR_120_-RXRα_98_-hSHP-1 complex (Fig. S4A). The XL-MS analysis of the SRC1_630-987_ and FXR_120_-RXRα_98_ complex resulted in the identification of 7 cross-links with high confidence (score ≥ 10) (Fig. 5D). Among these cross-links, three were observed between SRC1_630-987_ and RXRα_98_ (Fig. 5D). However, it was observed that the amino acids cross-linked to SRC1_630-987_ were all located outside the RXRα coactivator binding surface (Fig. 5E). This suggests that SRC1_630-987_ is not bound to the coactivator binding surface of RXRα-LBD. The remaining four cross-linking sites identified by XL-MS were located between SRC1_630-987_ and FXR_120_(Fig. 5D). Three of these pairs were found to be located on the coactivator binding surface of FXR-LBD and the NR-box3 of SRC1_630-987_. The other pair was also located near the coactivator binding surface of FXR-LBD (Fig. 5E). This suggests that NR-box3 of SRC1 is in close proximity to the coactivator binding surface of FXR. These findings imply that both NR-box2 and NR-box3 of SRC1 are involved in mediating the function and interaction of SRC1 with the FXR-RXRα heterodimer.

## Discussions

Given the significant therapeutic potential of FXR in treating MASH(41, 42), it is essential to comprehend the molecular mechanisms underlying FXR transcriptional regulation to aid in the rational drug design. In this study, we generate the first model for the complex formed by agonist/DNA-bound FXR-RXRα heterodimer and the complete SRC1-NRID domain. Using this model, we propose a mechanism elucidating how agonists and DNA co-regulate the recruitment of SRC1 by FXR-RXRα (Fig. 6).

**Figure 6.**
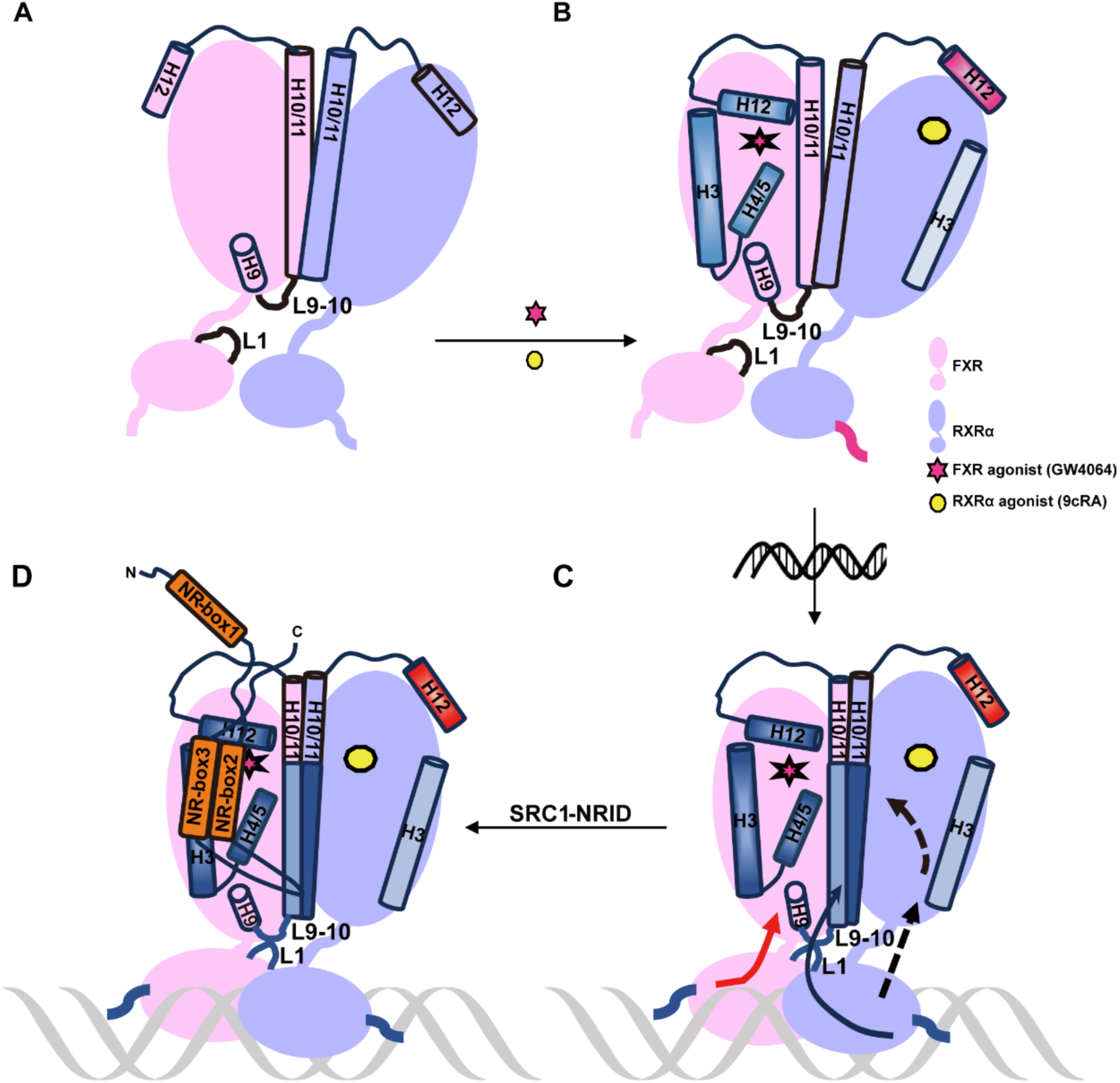
Model of agonist and DNA co-regulation of FXR-RXRα recruiting SRC1. (A) Model of FXR-RXRα without agonist binding. (B) Schematic of FXR-RXRα binding with agonists. Upon agonist binding to FXR-RXRα (red stars for FXR agonist GW4064; yellow circles for RXRα agonist 9cRA), the cofactor binding surfaces of FXR (H3, H4, and H12) is stabilized, facilitating coactivator recruitment. In contrast, the H12 of RXRα-LBD becomes more flexible upon agonist binding, impeding coactivator recruitment (blue indicates structural stability; red indicates flexibility, with color intensity showing the degree of stability or flexibility). The DBD of RXRα also becomes more flexible upon agonist binding, potentially aiding its binding to DNA response elements. (C) Schematic of FXR-RXRα when binding agonist and FXRE. FXRE binding signals are transmitted at FXR and RXRα, respectively. On one hand, the binding of FXRE compacts the structure of FXR-RXRα, bringing FXR-LBD and FXR-DBD closer together. The L9-10 of FXR-LBD and the L1 of RXRα-DBD transmit signals from the DNA to the FXR-DBD and then to the FXR-LBD, further stabilizing the coactivator binding surface (red arrow). On the other hand, FXR and RXRα transmits information through its DBD, enhancing the dimerization of the LBD of FXR and RXRα through the allosteric effect of RXRα (black arrow). This joint action promotes the stabilization of the coactivator binding region on the surface of FXR-LBD, which is conducive to the recruitment of coactivators. (D) Schematic representation of the asymmetric recruitment of SRC1 to FXR-RXRα-FXRE. NR-box refers to the α-helix formed by the LXXLL motif.

Interestingly, an interaction interface formed by the L9-10 of FXR-LBD and the L1 of FXR-DBD was found in our model, and this interface was important for the transcriptional activation activity of FXR-RXRα (Fig. 2). The similar interface has been observed in other solved crystal structures of NR multi-domain protein and DNA complexes(43), such as PPARγ-RXR (PDB: 3DZU), LXR-RXR (PDB: 4NQA), RAR-RXR (PDB: 5UAN). Our findings, for the first time, reveal a novel insight that DNA binding is the key factor that brings the DBD and LBD of FXR closer together, facilitating the formation of DBD-LBD interface (Fig. S2). We also observed that DNA binding enhances the heterodimerization and promote FXR coactivator recruitment (Fig. 6C).

Our HDX-MS studies have revealed that agonists binding to FXR-RXR heterodimer not only protects the LBP of FXR but also stabilizes its coactivator binding surface. Whereas agonists binding increases the flexibility of RXRα-DBD within the heterodimer, a similar phenomenon was also observed in VDR-RXR(18). The increased RXR-DBD flexibility after ligands binding may be a common feature among NR heterodimers, which can facilitate rapid recognition of specific DNA sequences.

In contrast to the crystal structure using only short peptides, our biochemical and cellular findings demonstrate that only one SRC1, containing the complete NRID, asymmetrically binds to FXR in FXR-RXRα through its NR-box2 and box3 (Fig. 6D). We note that in the crystal structure of PGC-1α (with the complete NRID domain) bound to HNF4α-LBD, only one LXXLL motif is observed, but electron density analysis implies the potential presence of multiple LXXLL motifs(44). Previously, one crystal structure of FXR-LBD (PDB ID: 1OSV) showed its capability to interact with two LXXLL peptides, with one peptide in the conventional coactivator binding region and the other in the region directly next to the first(11). These results imply that the interaction between NRs and coactivators is intricate, and the coactivator binding interface may not exclusively recruit coactivators through a single LXXLL motif.

Furthermore, we note that the pattern of asymmetric coactivator binding to one monomer of the RXR heterodimer was also observed in RAR-RXR, VDR-RXR, PPARγ-RXRα, and TR-RXRα(45, 46), and even between PGC-1α and ERRα/ERRγ homodimers(47). Although studies have demonstrated that each monomer in the CAR-RXR heterodimer can recruit SRC1, it is important to highlight that the protein used in these studies only consisted of the LBD domain(48). Our findings showed that DBD and DNA have a significant impact on the binding of coactivators to NR (Fig. 1C). Studying multi-domain NR can provide more physiologically relevant results. The asymmetric binding of coactivators to NR dimers we proposed here for FXR may suggest a common feature for NR. However, the molecular mechanisms underlying this phenomenon are still not fully understood. Our HDX-MS results demonstrated that agonists binding increase flexibility of H12 in RXRα-LBD, and this effect is further enhanced by DNA binding. The increased flexibility in H12 within one monomer of the NR dimer could potentially explain the widespread asymmetric binding of coactivators to the NR dimer.

Studies by Maletta et al. have shown that different DNA can induce different conformations in NR, directly affecting the spatial arrangement of NR cofactors(49). And there is a correlation between the asymmetry of the NR complex structure and the spatial localization of proteins involved in chromosome remodeling, as this localization could potentially impact the transcriptional activity of downstream genes(45, 49). So, we speculate that in ligand-dependent activated nuclear receptors, conformational changes induced by specific small molecule binding facilitate the binding of DBD to specific DNA motifs, followed by recruitment of coactivators and coregulators. The specific combinations of NRs, coactivators, and coregulators, as well as their positioning and interactions within the gene regulatory network, confer a highly specific impact on the transcriptional activity of genes. This will provide valuable insights for the design of gene-selective agonists.

Our work not only enhances the comprehension of the structure and function of the FXR transcriptional regulatory complex, but also adds to the increasing body of evidence highlighting the intricate nature of coactivator binding to NR. This will have important implications for the development of treatment strategies for related diseases.

## Materials and Methods

### Protein expression and purification

The FXR_120-486_ and RXRα_98-462_ were cloned into the pET-Duet-1 vector. Plasmids were transformed into E. coli Rosetta2 (DE3) cells and cultured in lysogeny broth (LB) medium at 37 ℃ until the optical density at 600 nm reached 0.6-0.8. Recombinant protein was induced by the addition of 0.1 mM isopropyl β-D-1-Thiogalactopyranoside (IPTG) and incubated at 16 ℃ for 20 hours. Cells were harvested by centrifugation at 4000 g for 10 minutes, lysed in buffer containing 20 mM Tris pH8.0, 300 mM NaCl, 1 mM TCEP, and 1 mM PMSF, and then subjected to ultracentrifugation at 13400 g for 40 minutes at 4 ℃. After centrifugation, the supernatant was loaded onto a Ni-NTA agarose resin, and proteins were eluted with the addition of 300 mM imidazole. The FXR and RXRα heterodimer were further purified using a Source Q anion exchange column and a gel filtration column. The final elution buffer for the Superdex 200 10/300 GL column (GE Healthcare) was 20 mM HEPES pH 7.5, 150 mM NaCl, and 0.5 mM TCEP.

To investigate the truncation containing three LXXLL motifs within the SRC1 protein (630–987), the corresponding fragment was cloned into the pET28a vector. The plasmids were transformed and induced, and cells were collected as previously described. The cells were lysed in a solution of 20mM Tris pH 7.5, 300 mM NaCl, and 6 mM β-Mercaptoethanol (β-Me), and the supernatant was purified by Ni-NTA column. Subsequently, the SRC1_630-987_ protein was further purified using Source Q and gel filtration chromatography columns according to the previously described protocol.

The human SHP-1 sequence (5′-GATCCCTGAGTTAATGACCTTGT-3′) and its corresponding antisense strand were synthesized by Azenta Life Science. The two strands were solubilized in a buffer containing 20 mM HEPES, pH 7.5, 150 mM NaCl, and then diluted to a concentration of 1 mM. Subsequently, the two strands were annealed by combining equal amounts of each strand and heating them to 95°C, followed by a slow cooling to 25°C.

To purify SRC1_630-987_-FXR_120_-RXRα_98_-hSHP-1, purified SRC1_630-987_ and FXR_120_-RXRα_98_ were combined at a molar ratio of 1:4, along with the FXR agonist GW4064 and RXRα agonist 9cRA. Subsequently, hSHP-1 was added to the mixture at a molar ratio of 1:1.5 relative to FXR_120_-RXRα_98_. The mixture was thoroughly mixed and incubated on ice for 2 hours. Afterward, it was centrifuged at 12000 rpm for 10 minutes at 4 °C, followed by purification using Superdex 200 10/300 GL with a buffer containing 20 mM HEPES pH 7.5, 150 mM NaCl, and 0.5 mM TCEP.

### BLI Kinetic Assay

The binding affinities of SRC1_630-987_ and hSHP-1 towards FXR_120_-RXRα_98_ were determined using BLI on an Octet R8 (Sartorius). All experiments were performed at 25 °C in a buffer containing 20 mM HEPES pH 7.5, 150 mM NaCl, 0.02% Tween-20. Purified 6×His-tagged FXR-RXRα protein was diluted with HEPES buffer into different concentrations (i.e., 1000 nM, 500 nM, 250 nM, 125 nM). Streptavidin biosensors were pre-equilibrated in the buffer for at least 10 min before use in experiments. The biotinylated SRC1 and hSHP-1 proteins were loaded onto the streptavidin biosensors for durations of 60 s and 40 s, respectively. Following a period of association for 210 s, the biosensors were transferred to the HEPES buffer to measure dissociation over a period of 420 s. Results were analyzed by ForteBio Data Analysis software.

### Cross-linking mass spectrometry

Crosslinking reactions were conducted in 100 µL protein solutions in 20 mM HEPES pH 7.5, 150 mM NaCl, 1 mM TCEP. FXR-RXRα samples with and without ligands and SRC1_630-987_ were crosslinked in HEPE buffer using Bis[sulfosuccinimidyl] suberate (BS3, ThermoFisher) and 1-(3-Dimethylaminopropyl)-3-ethylcarbodiimide hydro (EDC, ThermoFisher). For BS3 crosslinking, freshly prepared 12.5 mM stock solution of BS3 in sterilized distilled water were added in 30-, 50- and 70-fold molar ratio to protein samples. Crosslinking reactions were conducted during 30 min at room temperature and further quenched during 15 min using Tris pH7.5 to a final concentration of 50 mM. For EDC crosslinking, freshly prepared 60 mM solution of EDC were added in 30-50- and 70-fold molar ratio to protein samples. Then activator Sulfo-NHS was adding 2.5-fold molar ratio to EDC, mixed evenly, and reacted at room temperature for 2 hours or overnight on ice. The cross-linking reaction was terminated using a final concentration of 50 mM Tris pH 7.5 and 20 mM β-Me at room temperature for 15 min.

The gel band stained with Coomassie Brilliant Blue was fragmented and subjected to a series of washing steps, including water, 50 mM NH_4_HCO_3_ in 50% acetonitrile, and 100% acetonitrile. Following this, the protein was treated with 10 mM TCEP (ThermoFisher Scientific) in 100 mM NH_4_HCO_3_ at room temperature for 30 minutes for reduction and then alkylated with 55 mM iodoacetamide (Sigma) in 100 mM NH_4_HCO_3_ in the dark for 30 minutes. Subsequently, the gel pieces were washed with 100 mM NH_4_HCO_3_ and 100% acetonitrile, followed by drying with a SpeedVac. The dried fragments were then digested with trypsin by adding 12.5 ng/μL trypsin (Promega) in 50 mM NH_4_HCO_3_ for 16 hours at 37°C. The resulting tryptic peptides were extracted twice with 50% acetonitrile/5% formic acid and dried using a SpeedVac. The sample was reconstituted with 0.1% formic acid, desalted with a MonoSpin^TM^ C18 column (GL Science, Tokyo, Japan), and dried once more with a SpeedVac.

Utilizing a custom-made 30 cm-long pulled-tip analytical column (75 μm ID containing ReproSil-Pur C18-AQ 1.9 μm resin from Dr. Maisch GmbH, Germany), the peptide mixture underwent analysis. Subsequently, the column was integrated with an Easy-nLC 1200 nano HPLC (ThermoFisher Scientific, San Jose, CA) for mass spectrometry investigation. Maintaining a temperature of 55 ℃, the analytical column was used throughout the experiments.

Analysis of MS/MS data was carried out using a Q Exactive Orbitrap mass spectrometer (ThermoFisher Scientific). Control of MS scan functions and LC solvent gradients was managed by the Xcalibur data system provided by Thermo Scientific.

The identification of peptides with isopeptide bonds was performed using the pLink2 software (pFind Team, Beijing, China) according to previously established methods (50, 51). Parameters for the pLink search included: enzyme: trypsin; maximum missed cleavages: 3; tolerance for precursor and fragment ions: 20 ppm. Carbamidomethylation of cysteine was specified as a static modification, while oxidation of methionine was considered as a dynamic modification. The final results were refined using a 5% false discovery rate threshold at the spectral level.

### Small-Angel X-ray scattering (SAXS) data collection and processing

SAXS data is collected by SEC-SAXS on the BL19U2 beamline station of National Facility for Protein Science in Shanghai (NFPS) and Shanghai Synchronous Radiation Facilities (SSRF). The Superdex 200 Increase 10/300 GL column was equilibrated with a buffer solution comprising 20 mM Tris-HCl pH 7.5, 150 mM NaCl and 0.5 mM TCEP. Subsequently, a 15 mg/ml solution of FXR_120_-RXRα_98_-FXRE was loaded onto the column. The data collection wavelength was set at 1.033 Å. Use BioXTAS RAW software(52) to take the average value of the scattering data, and then convert the 2D scattering image to a 1D SAXS curve. Use the GNOM(53) in the ATSAS software package to calculate the particles P(r) and the maximum sizes D max based on the 1D SAXS curve, and use the DAMMIF to determine Low-resolution shapes from solution scattering data(54, 55). The fitting model’s quality was assessed by computing the χ^2^ value using CRYSOL(53), and the fit of the docking model to the Ab initio modeling model computed from the experimental SAXS data is done by CIFSUP.

### Size-Exclusion Chromatography Coupled to Multi-Angle Light Scattering (SEC-MALS)

The absolute molecular weights of protein complexes in solution were determined using size exclusion chromatography (SEC) coupled with multi-angle light scattering detection (SEC-MALS).

Samples at a concentration of 3 mg/ml were loaded onto GE Healthcare Superdex 200 10/300 GL pre-equilibrated using buffer containing 20 mM HEPES pH 7.5, 150 mM NaCl and 1 mM TCEP. The samples separation and subsequent detection were performed using an Agilent HPLC system and a MALS instrument (DAWN+ECLIPSE; Wyatt Technologies, USA) at a flow rate of 0.5 ml/min. Data were processed using ASTRA 8 software.

### Structural modelling of FXR-RXRα binding to hSHP-1

Atomic models of FXR-RXRα-IR1 were constructed by leveraging crystal structures from FXR-RXRα-DBD (PDB: 5Z12,6A5Z) and FXR-RXRα-LBD-IR1 (PDB: 8HBM) using DISVIS(56, 57), and the LZerD Web server(39, 40) in combination with distance constraint information provided by XL-MS. Initially, crosslinking pairs’ data in the protein complexes were analyzed with DISVIS, eliminating pairs exceeding the crosslinker’s distance limit and those exhibiting unreasonable configurations. This analysis also helped identify potential active residues. Through DISVIS analysis of 26 high-intensity cross-linking peptides (pLINK2 calculated score ≥ 10), information on active residues on the surface interface between FXR-RXRα-LBD and FXR-RXRα-DBD-IR1 was extracted. Subsequently, utilizing this data, protein models were docked employing the LZerD web server, resulting in a series of composite models.

### Hydrogen/deuterium exchange (HDX) mass spectrometry experiments

All three samples (FXR_120_-RXRα_98_, FXR_120_(GW4064)-RXRα_98_(9cRA), FXR_120_(GW4064)-RXRα_98_(9cRA)-hSHP-1) were prepared in a buffer composed of 20 mM HEPES at pH 7.4 and 150 mM NaCl, resulting in a final complex concentration of 3 mg/ml per reaction. The automated processing of samples was facilitated by a LEAP Technologies Hydrogen Deuterium Exchange PAL system located in Carrboro, NC. Chromatographic separation was conducted using a U3000 RSLC nano HPLC system. Hydrogen Deuterium Exchange (HDX) measurements were recorded at various time points (0 s, 10 s, 30 s, 120 s, 600 s, and 1800s) while maintaining a temperature of 4 °C. In the chromatography process, mobile phases A and B consisted of 0.1% formic acid in H_2_O and 0.1% formic acid in an acetonitrile solution, respectively. Following deuteration with D_2_O, all samples were quenched using a buffer solution (200 mM citric acid, 4 M guanidine-HCl, 500 mM TCEP in H_2_O, pH 2.5). Subsequently, the samples were digested with pepsin, trapped on a C18 trap, and separated on C18 analytical chromatography columns. For peptide identification, a Thermo LTQ Orbitrap-Elite mass spectrometer (San Jose, CA) was used for detection and analysis. HDX experiments were realized in triplicate for each time point. Mass spectra was adopted data-dependent acquisition (DDA) mode at the *m/z* range of 300-1500 to provide Peptide Source for data analysis. The spectra generated were searched in PEAKS online with a home-made protein sequence library to screen Peptide Source. Retention time and sequence information for each peptide was carried out using HDExaminer 2.0 (Sierra Analytics Inc., Modesto, CA). Using DDA data as reference, deuterium uptake of peptides and predicted amino acids was calculated. The uptake of deuterium was calculated using the software algorithm via matching the best theoretical isotope distribution pattern to the observed isotope distribution pattern. Statistical significance for the differential HDX data is determined by t-test for each time point.

### Luciferase assay

Luciferase reporter gene assays were conducted using HEK293T cells. The cells were seeded at a density of 25,000 cells/well in a 96-well plate and cultured in DMEM supplemented with 10% fetal bovine serum at 37 °C, 5% CO_2_ for 24 hours. Co-transfection was performed using Lipofectamine 2000 (Thermo Fisher Scientific) and Opti-MEM, with a mixture containing 70 ng of hSHP-1-luciferase reporter gene, 70 ng of pcDNA3.1-human FXR, RXRα, and SRC1, and 10 ng of Renilla internal reference control. After 6-8 hours of transfection, the Opti-MEM medium was replaced with DMEM. Concurrently, ligands for nuclear receptors (GW4064 for FXR and 9cRA for RXRα) were added at a final concentration of 100 nM. After 24 hours of treatment, the cells were lysed, and the transcriptional activity was quantified using the dual luciferase assay system from Promega.

## Acknowledgements

We thank the staff from BL19U2 beamlines at National Facility for Protein Science in Shanghai (NFPS) and Shanghai Synchrotron Radiation Facility, for assistance during data collection. We thank the Mass Spectrometry System at the National Facility for Protein Science in Shanghai (NFPS), Shanghai Advanced Research Institute, Chinese Academy of Science, China for MS sample preparation, data collection and data analysis. We thank Baizhen Biotechnologies Inc. for HDX MS experiment, including data collection and analysis. We thank Nannan Wang at the Proteomics and Metabolomics Platform, Guangzhou Laboratory for help with SEC-MALS data collection and analysis.

The work was financially supported by grants from the National Key Research and Development Program (2022YFA13031001), National Natural Science Foundation of China (32301008).

## Author Contributions

Y.S., Y.G., M.S., Y.D., YY., Y.W., C.P. and Y. X. performed experiments. J.L, Y.S and N.W designed the experiments. J.L, Y.S and N.W wrote the manuscript.

## Supplementary materials

**Figure. S1.**
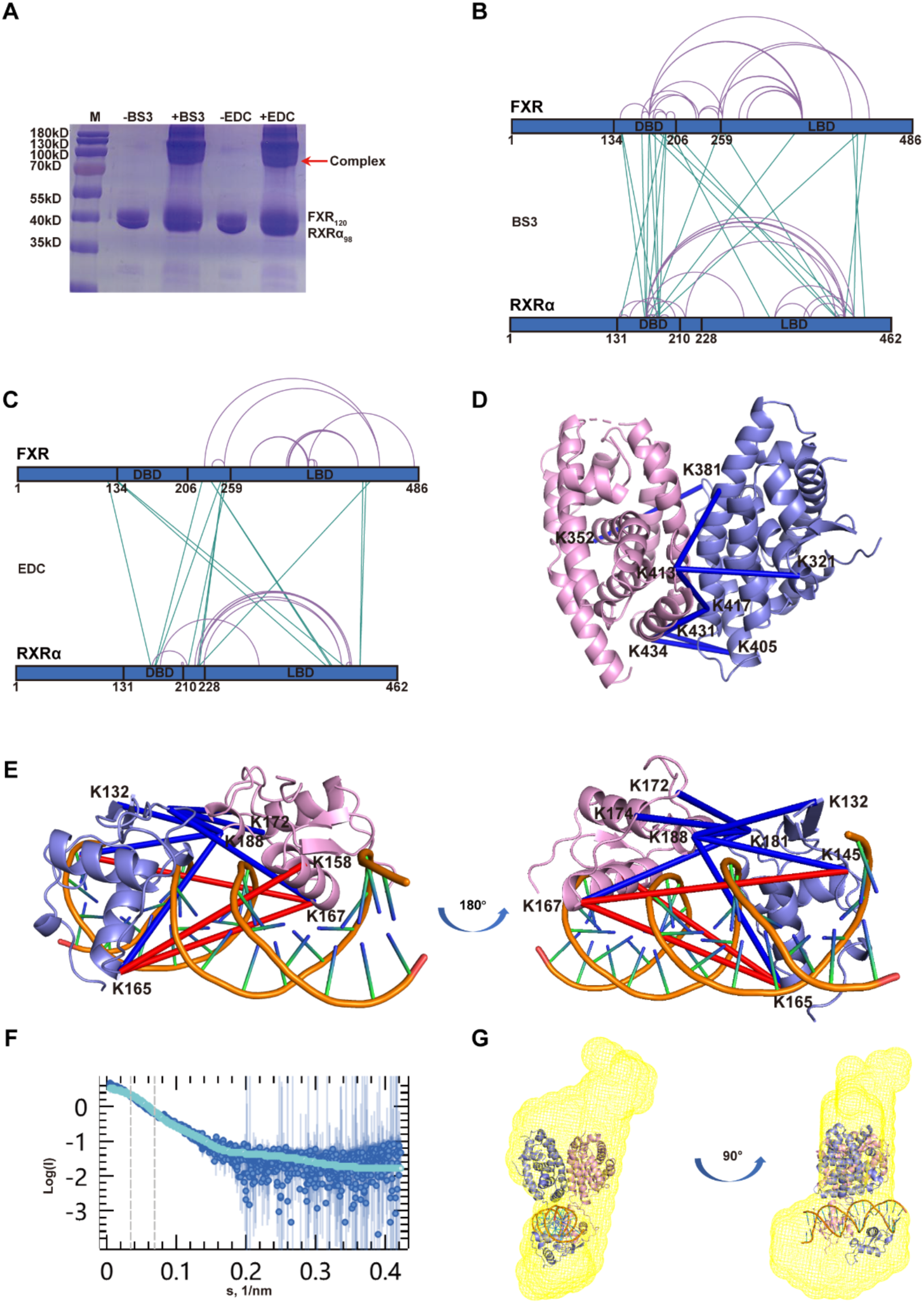
Integrative model of the FXR-RXRα heterodimer binding to hSHP-1. (A) SDS-PAGE analysis result of FXR_120_-RXRα_98_-hSHP-1 with and without cross-linker (BS3 and EDC). (B and C) The high-confidence BS3 and EDC XL-MS results for FXR_120_-RXRα_98_-hSHP-1 are illustrated on the sequences of FXR and RXRα. The figure displays XL-MS data with scores ≥ 10, with green lines indicating cross-links and purple lines indicating self-links. (D and E) High-confidence cross-links (score ≥ 10) between the LBD and DBD of FXR and RXRα are marked on the crystal structures of FXR-RXRα-LBD and FXR-RXRα-DBD (FXR in pink; RXRα in slate). Cross-links consistent with the structure are colored blue, while those not consistent with the structure are colored red. (F) The experimental SAXS profile is overlaid with the theoretical profile calculated from the modeled structure of FXR_120_-RXRα_98_-hSHP-1. The theoretical profile is depicted in light blue, while the experimental SAXS profile is shown in dark blue. (G) The structural model of FXR_120_-RXRα_98_-hSHP-1 is superimposed on the ab initio structure derived from SAXS. The cartoon represents the structural model of FXR_120_-RXRα_98_-hSHP-1, while the green mesh corresponds to the ab initio structure.

**Figure. S2.**
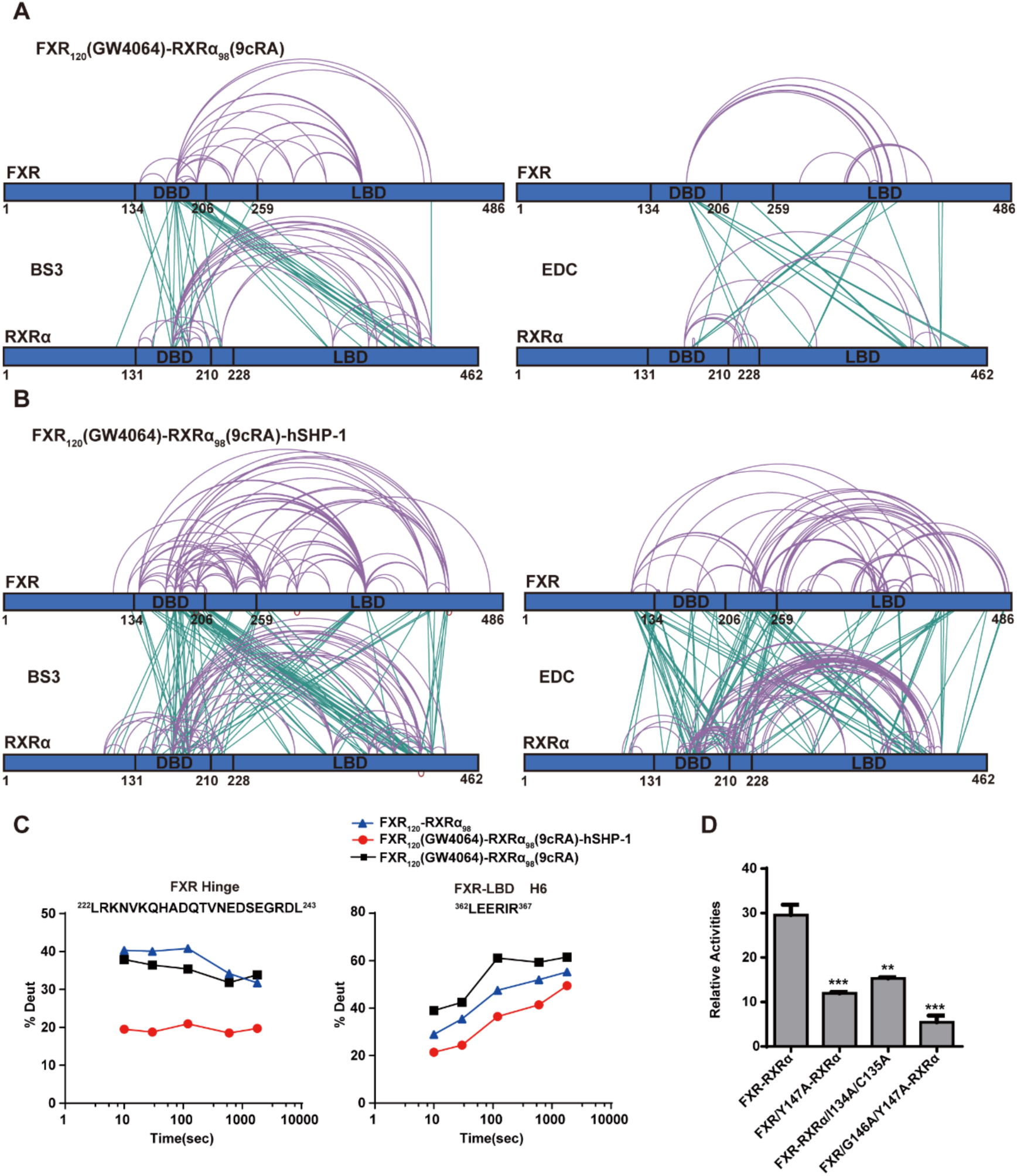
Allosteric communication between the LBD and DBD of FXR-RXRα. (A and B) Schematic maps illustrating BS3 or EDC cross-linked residues of FXR_120_(GW4064)-RXRα_98_(9cRA) and FXR_120_(GW4064)-RXRα_98_(9cRA)-hSHP-1 are presented on the FXR and RXRα sequences. Interactions between molecules are denoted by green lines, indicating intermolecular cross-linking, while intramolecular cross-linking is depicted by purple lines. (C) Deuterium uptake plot of the hinge region and H6 of FXR-LBD. (D) Luciferase reporter gene assay was used to detect the effect of mutations in the NTD of FXR and RXRα on the transcriptional activity of FXR_120_-RXRα_98_ on hSHP-1 in the presence of GW4064 and 9cRA.

**Figure. S3.**
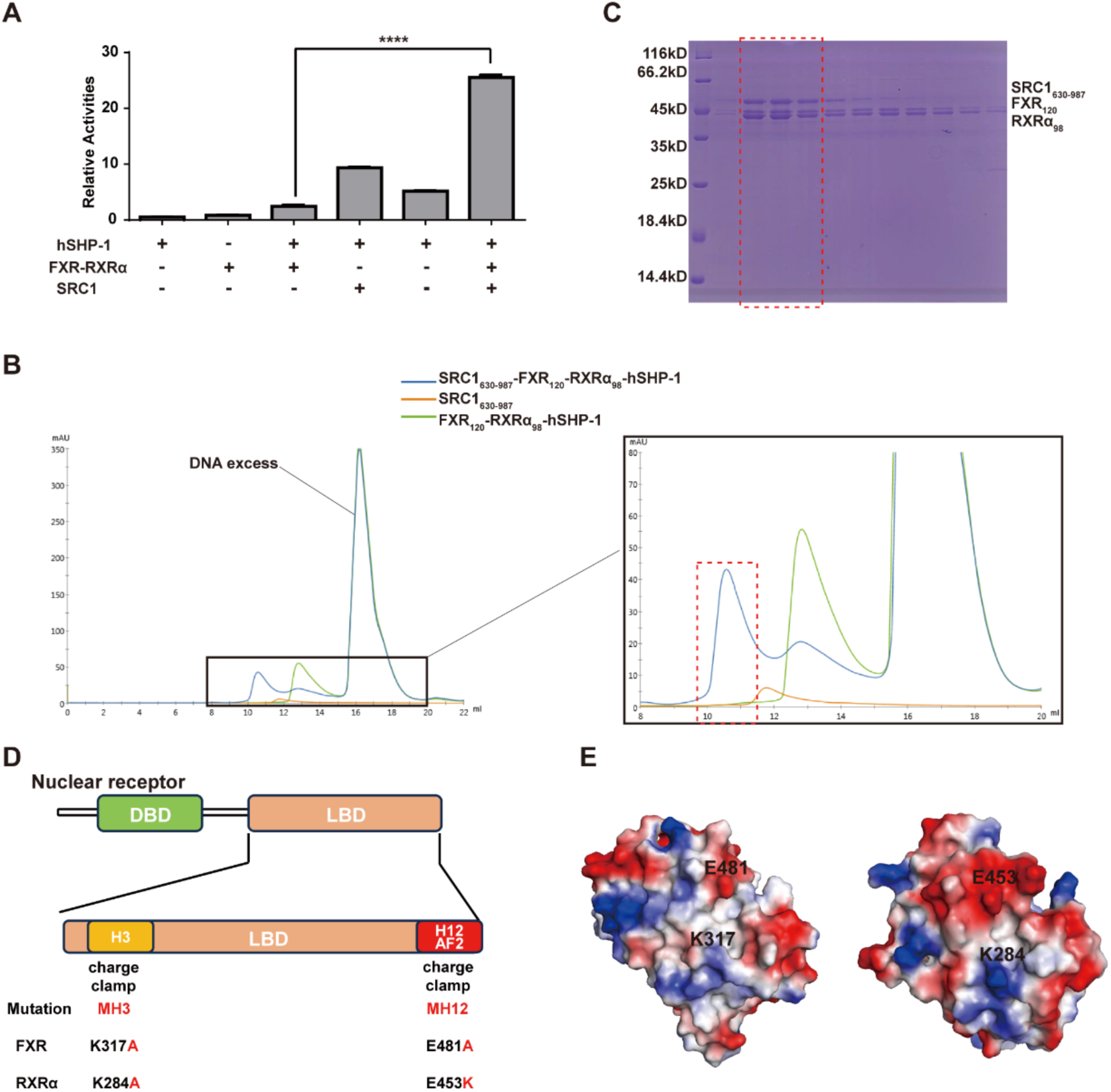
The transcriptional activation of FXR-RXRα is affected by the asymmetric binding of SRC1 to FXR. (A) The impact of SRC1 on the transcriptional activity of FXR-RXRα in stimulating hSHP-1 was examined through a luciferase reporter gene assay conducted in the presence of GW4064 and 9cRA. (B) Overlay of UV280 absorption peaks from SEC of FXR_120_-RXRα_98_-hSHP-1, SRC1_630-987_, and their complexes, with the region of interest magnified for better visualization. (C) SEC fractions were analyzed by SDS-PAGE, and the sample well corresponding to the framed peak in Figure C is highlighted by a red dotted line. (D) Schematic representation of the FXR and RXRα mutants with charge-clamp modifications used in this study. (E) Electrostatic potential surfaces corresponding to the FXR-LBD and RXRα-LBD. Positive potential regions are indicated in blue, negative potential regions in red, and neutral hydrophobic regions in white. Specifically, K317 and E481 constitute the ’charge clamp’ in FXR, while K284 and E453 constitute the ’charge clamp’ in RXRα.

**Figure. S4.**
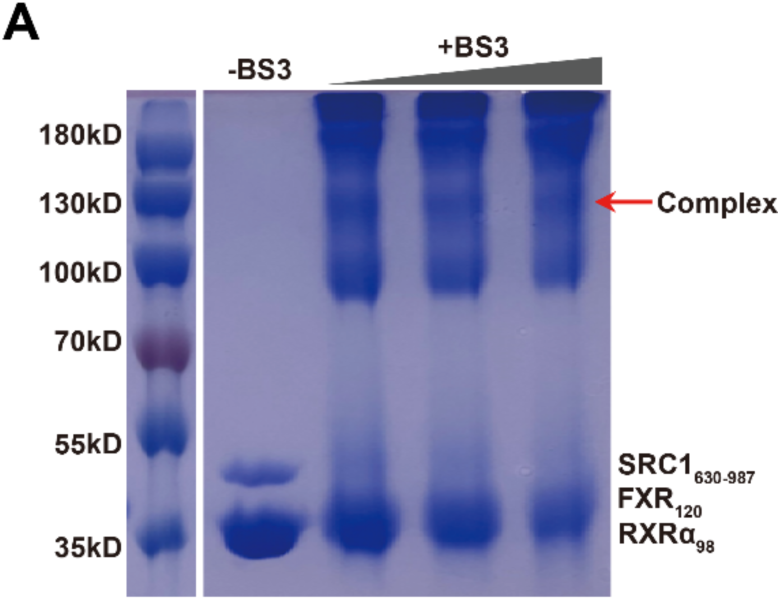
(A) SDS-PAGE results of SRC1_630-987_-FXR_120_-RXRα_98_-hSHP-1 samples with and without cross-linker (BS3). The band indicated by the red arrow corresponds to the band analyzed by mass spectrometry.

**Table S1.** SAXS sample details and parameters.

| Data Collection Parameters |  |  |  |  |
| --- | --- | --- | --- | --- |
| Sampel | | FXR<sub>120</sub>-RXRα<sub>98</sub>-hSHP-1 | SRC1<sub>630-987</sub> | SRC1<sub>630-987</sub>-FXR<sub>120</sub>-RXRα<sub>98</sub>-hSHP-1 |
| SEC-SAXS column |  | Superdex 200 10/300GL | Superdex 200 10/300GL | Superdex 200 10/300GL |
| Loading concentration |  | 10 mg/mL | 10 mg/mL | 15 mg/mL |
| Flow rate |  | 0.4 mL/min | 0.45 mL/min | 0.45 mL/min |
| Solvent |  | 20 mM Tris-HCl, pH7.5, 150 mM NaCl 0.5 mM TCEP | 20 mM HEPES-NaOH, pH7.5, 150 mM NaCl 0.5 mM TCEP | 20 mM HEPES-NaOH, pH7.5, 150 mM NaCl 0.5 mM TCEP |
| Sample-detector distance (m) |  | 2.91 | 2.91 | 2.85 |
| Wavelength (Å) |  | 1.03 | 1.03 | 1.03 |
| Exposure time (s) |  | 1 | 1 | 1.5 |
| Temperature (°C) |  | 25 | 25 | 25 |
| Structural Parameters |  |  |  |  |
| From Guinier fit | I (0) | 3.92±0.03 | 4.05±0.04 | 14.65±0.13 |
|  | Rg (Å) | 41.31±0.45 | 53.98±0.83 | 65.24±0.72 |
| From *P*(*r*) | I (0) | 3.94±0.03 | 4.18±0.03 | 14.73±0.01 |
|  | Rg (Å) | 42.8±0.42 | 59.63±0.66 | 68.90±0.56 |
| Molecular Mass Determination |  |  |  |  |
| MM (kDa) from V(c) |  | 79.10 | 34.9 | 130.00 |
| Protein sequence MW (kDa) |  | 85.10 | 39.01 | 124.11 |
| Modeling |  |  |  |  |
| DAMMIF | NSD | 0.89±0.15 | 0.94±0.08 | 0.80±0.09 |
|  | Ensemble Resolution | 38.23 | 59.56 | 62.91 |

